# First brain de-novo transcriptome of Tyrrenhian tree frog, *Hyla sarda*, for the study of dispersal-related behavioral variation

**DOI:** 10.1101/2022.05.25.493414

**Authors:** Pietro Libro, Roberta Bisconti, Andrea Chiocchio, Giada Spadavecchia, Tiziana Castrignanò, Daniele Canestrelli

## Abstract

Dispersal is a key process in ecology and evolution as it contributes to shaping the spatial patterns of biological diversity at all its levels of organization. Growing evidence is unveiling the role of phenotypic trait variation in affecting dispersal dynamics of populations, and that substantial, rapid, and directional changes in the phenotypic makeup of populations would occur during range expansions by spatial sorting of dispersal-related traits. Accordingly, selective pressures at the front of a range expansion wave would actively promote individuals with higher dispersal abilities. Although many studies have been focusing on phenotypic trait evolution during range expansion, its genomic underpinnings are almost unexplored, hampering a thorough understanding of the evolutionary processes involved during dispersal.

The Tyrrhenian tree frog Hyla sarda is a small amphibian endemic to the Tyrrhenian islands. According to previous phylogeographic studies, H. sarda underwent a northward range expansion from the north of Sardinia (western Mediterranean sea) during the last glacial phase. The colonization of the Corsica island was allowed by the temporary formation of a wide land bridge connecting Sardinia and Corsica, induced by the marine regression occurred during the last glacial maximum. The postglacial loss of this land bridge prevented any subsequent gene flow between Corsican and Sardinian populations. As a consequence, the genetic and phenotypic legacies of this range expansion event can still be detected in current populations, which makes H. sarda an interesting candidate species for the study of the genetic underpinnings of phenotypic trait evolution during range expansions.

Here, we contribute to the investigation of the genetic underpinning phenotypic trait evolution during range expansion, by generating the first brain de-novo transcriptome of H. sarda. We focused on the brain transcriptome as behavioural variation at personality traits has been identified as a key component of the dispersal syndromes, substantial personality traits variation has been observed within H. sarda populations, and brain gene expression patterns have been linked to a vast number of behavioural responses to environmental stimuli.

## Background & Summary

Dispersal is a key process in ecology and evolution as it contributes to shaping the spatial patterns of biological diversity at all its levels of organization (Kokko and López–Sepulcre, 2006; Clobert et al., 2012). Growing evidence is unveiling the role of phenotypic trait variation in affecting dispersal dynamics of populations (Liedvogel et al., 2011), and that substantial, rapid, and directional changes in the phenotypic makeup of populations would occur during range expansions by spatial sorting of dispersal-related traits (Canestrelli et al., 2016a,b; Miller et al., 2020). Accordingly, selective pressures at the front of a range expansion wave would actively promote individuals with higher dispersal abilities (Lindström et al., 2013; Canestrelli et al., 2016a, b; Pizzatto et al., 2017). Although many studies have been focusing on phenotypic trait evolution during range expansion, its genomic underpinnings are almost unexplored, hampering a thorough understanding of the evolutionary processes involved during dispersal.

The Tyrrhenian tree frog *Hyla sarda* (De Betta, 1853) is a small amphibian endemic to the Tyrrhenian islands. According to previous phylogeographic studies, *H. sarda* underwent a northward range expansion from the north of Sardinia (western Mediterranean sea) during the last glacial phase (Spadavecchia et al., 2021). The colonization of the Corsica island was allowed by the temporary formation of a wide land bridge connecting Sardinia and Corsica, induced by the marine regression occurred during the last glacial maximum (Bisconti et al., 2011a, b; Spadavecchia et al., 2021). The postglacial loss of this land bridge prevented any subsequent gene flow between Corsican and Sardinian populations. As a consequence, the genetic and phenotypic legacies of this range expansion event can still be detected in current populations (Liparoto et al., 2020; Canestrelli et al., 2021; Bisconti et al. 2022), which makes *H. sarda* an interesting candidate species for the study of the genetic underpinnings of phenotypic trait evolution during range expansions.

Here, we contribute to the investigation of the genetic underpinning phenotypic trait evolution during range expansion, by generating the first brain *de-novo* transcriptome of *H. sarda*. We focused on the brain transcriptome as behavioural variation at personality traits has been identified as a key component of the dispersal syndromes (Canestrelli et al., 2016a), substantial personality traits variation has been observed within *H. sarda* populations (Bisconti et al. 2022), and brain gene expression patterns have been linked to a vast number of behavioural responses to environmental stimuli (Whitfield et al., 2003; Bendesky and Bargmann, 2011; Rey et al., 2013; Harris & Hofmann, 2014; Bell et al., 2016).

## Methods

### Generation of the Datasets

Sampling procedures were performed under the approval of the Institute for Environmental Protection and Research ‘ISPRA’ (protocol # 5944), Ministry of Environment ‘MATTM’ (protocol #8275), Regione Sardegna (#12144) and Corsica (#2A20180206002 and #2B20180206001). Permission to temporarily house amphibians was granted by Local Health and Veterinary Centre, with license code 050VT427. All handling procedures outlined in the present study were approved by the Ethical Committee of the University of Tuscia for the use of live animals (D.R. n. 677/16 and D.R.644/17). We sequenced 9 Tyrrhenian tree frog individuals showing behavioral traits differentiation along the bold-shy axis of personality (Table 1). Sampling procedures and phenotyping were described in Bisconti et al. (2022). Brains were dissected and stored in RNAprotect Tissue Reagent (Quiagen). The RNeasy Plus Kit (Quiagen) was used to extract the total RNA, following the manufacturer instructions. RNA quality and concentration were evaluated by using both the spectrophotometer Agilent Cary60 UV-vis and the Bioanalyzer Agilent 2100 (Agilent Technologies, Santa Clara, USA).

**Table 1.** Summary of the 9 samples and library deposited in the ENA (European Nucleotide Archive, Study Accession Id PRJEB51203), in terms of number of raw and trimmed reads per sample.

| Sample code | Sample location | Phenotype | Run ID | Raw sequences | Trimmed sequences (% of total reads) |
| --- | --- | --- | --- | --- | --- |
| HS 9 | Porto Pollo, Sardinia | Bold | ERR9486926 | 72098178 | 69503761 (96.4) |
| HS 17 | Porto Pollo, Sardinia | Bold | ERR9486869 | 53786371 | 52007828 (96.7) |
| HS 27 | Cala Ginepro, Sardinia | Bold | ERR9486870 | 54282464 | 52269332 (96.3) |
| HS 44 | Cala Ginepro, Sardinia | Bold | ERR9486863 | 49300066 | 47606858 (96.6) |
| HS 56 | Elba | Bold | ERR9433479 | 51227766 | 46626281 (91.0) |
| HS 87 | Etagn de Canettu, Corsica | Shy | ERR9433456 | 53775786 | 50544832 (94.0) |
| HS 93 | Etagn de Canettu, Corsica | Shy | ERR9433396 | 54008324 | 50503363 (93.5) |
| HS 104 | Aleria, Corsica | Shy | ERR9433394 | 51829828 | 47628168 (91.9) |
| HS 111 | Aleria, Corsica | Shy | ERR9433393 | 52754498 | 49148365 (93.2) |

Library preparation and RNA sequencing were performed by NOVOGENE (UK) COMPANY LIMITED using the Illumina NovaSeq platform. Library construction was carried out by using the NEBNext® Ultra ™ RNA Library Prep Kit for Illumina®, according to the manufacturer instructions. Briefly, the mRNA present in the total RNA sample was isolated by means of magnetic beads made of oligos d(T)25 (i.e., polyA-tailed mRNA enrichment). Subsequently, mRNA was randomly fragmented, and the cDNA was synthesized using random hexamers and the reverse transcriptase enzyme. Once the synthesis of the first chain has finished, the second chain was synthesized with the addition of an Illumina buffer, dNTPs, RNase H and polymerase I of *E*.*coli*, by using the Nick translation method. The resulting products went through purification, repair, A-tailing and adapter ligation. Finally, the fragments of the appropriate size were enriched by PCR, the indexed P5 and P7 primers were introduced, and the final products were purified.

The obtained libraries were sequenced using the Illumina Novaseq6000 sequencing system, through a paired-end 150bp (PE150) strategy. We obtained on average 54.78 million reads for each library. The sequencing data are available at the European Nucleotide Archive (ENA, study ID PRJEB51203).

### Data Processing and Transcriptome Analyses

High-quality RNA-Seq reads (sequences) were used in the de-novo transcriptome assembly. All the described bioinformatics procedures were performed on the high performance computing system provided by ELIXIR-IT HPC@CINECA (Castrignanò et al. 2020).

Raw RNA-seq reads were pre-processed by removing adapters and low-quality sequences (<Q30) using Trimmomatic (v. 0.39) (Bolger et al. 2014) with default parameters. After the removal of the low-quality reads, 456,838,788 clean reads (i.e., 93% of raw reads) were maintained for building the de novo transcriptome assembly. Sequencing summary statistics showing the total number of reads before and after trimming and quality filtering are presented in Table 1. RNA-Seq read quality before and after trimming was assessed using FastQC (https://www.bioinformatics.babra-ham.ac.uk/projects/fastqc/) and aggregated using MultiQC (Ewels et al., 2016), read quality after trimming is presented in Supplementary Figure 1.

As there is no reference genome for *Hyla sarda*, two workflows were examined to choose an optimal transcriptome assembly strategy (Supplementary Figure 2). In order to construct an optimized de novo transcriptome avoiding chimeric transcripts, we adopted the strategy to launch two de novo assembly tools using a multi-kmer approach. The two assemblers, Trinity (trinityrnaseq-v2.11.0), and SPAdes (version 3.11.1) used in rnaSPAdes mode (Bushmanova et al., 2019), were launched to improve the reliability of the final assembly. The results from these analyzes, using both assemblers, are shown in Table 1s.

The outputs obtained by the assemblers were subjected to a validation step (“Transcriptome Quality Check” in (Supplementary Figure 2). In this case the software used are Busco (v. 4.1.4) (Simão et al., 2015) and TransRate (v. 1.0.3) (Smith-Unna et al., 2016). These tools generate several metrics that serve as a guide to evaluate error sources in the assembly process and provide evidence about the quality of the assembled transcriptome. TransRate is a contig validation software using four alignment linked quality measures to generate a global quality criterion for each contig and for the complete set. BUSCO, Benchmarking Universal Single-Copy Orthologs, provides a quantitative measure of transcriptome quality and completeness, based on evolutionarily informed expectations of gene content from the near-universal, ultra-conserved eukaryotic proteins (eukaryota_odb9) database (Supplementary Table 1).

The outputs obtained after validation were merged with Trans-ABySS (v2.0.1). After, the assembly result was the input for CD-HIT-est program (v4.8.1 used with default parameters, corresponding to a similarity 95%), (Fu et al., 2012) a hierarchical clustering tool used to avoid redundant transcripts and fragmented assemblies common in the process of de novo assembly, providing unique genes. CD-HIT-est orders the contigs by length and removes all the included ones given identity and coverage thresholds. The results are shown in Supplementary Table 1. TransDecoder (Tang et al., 2015), the current standard tool that identifies long open read frames (ORFs) in assembled transcripts, was run with predefined parameters on CD-HIT-est output file. TransDecoder by default performs ORF prediction on both strands of assembled transcripts regardless of the sequenced library. It also ranks ORFs based on their completeness, and to determine if the 5 ‘end is incomplete, looks for any length of AA codons upstream of a start codon (M) without a stop codon. Adopt the “Longest ORF” rule and select the highest 5 AUG (relative to the inframe stop codon) as the translation start site. Finally, we identified ORFs with homology to known proteins using BLAST search (Uniprot database, BLAST version 2.2.26+, BLASTp, with an e-value cut-off ≤ 10-5) and searched for protein signatures in the Pfam-A database. In the last step, the program TransDecoderPredict uses this information to predict the coding sequences.

#### Transcriptome annotation

We provided different annotations for all further analysis. Contigs were aligned with DIAMOND on NCBI NR, Swiss-Prot and TrEMBL to retrieve corresponding best annotations (Buchfink et al., 2015). An annotation matrix was then generated by selecting the best hit for each database. Following the analysis of blastx against NCBI Nr, Swiss-Prot and TrEMBL, we obtain respectively: 179944 sequences equal to (46.99%), 133455 (34.85%), 179978 (47.00%). The results obtained following the analysis with blastp against NCBI Nr, Swiss-Prot and TrEMBL are: 128401 (33.53%), 88996 (23.24%) and 130165 (33.99%). Information on the datasets resulting from this study is available in Table 2s. A Venn diagram was created to show the redundancy of the annotations in different databases; the diagrams have been constructed for both Diamond blastx (Supplementary Figure 3a) and Diamond blastp (Supplementary Figure 3b) results showing 132635 and 88671 unigenes commonly annotated from the three databases. In a second step, contigs were also processed with InterProScan (Hunter S, et al 2009) to scan InterProScan signatures. The InterPro database (http://www.ebi.ac.uk/interpro/) integrates together predictive models or ‘signatures’ representing protein domains, families and functional sites from multiple, diverse source databases.

#### Technical Validation

In Figure 1, the values of *raw reads, paired-reads* after trimming and *trimmed paired-reads* mapped against *Hyla sarda* de novo transcriptome are expressed as million and always higher than 90%.

**Figure 1.**
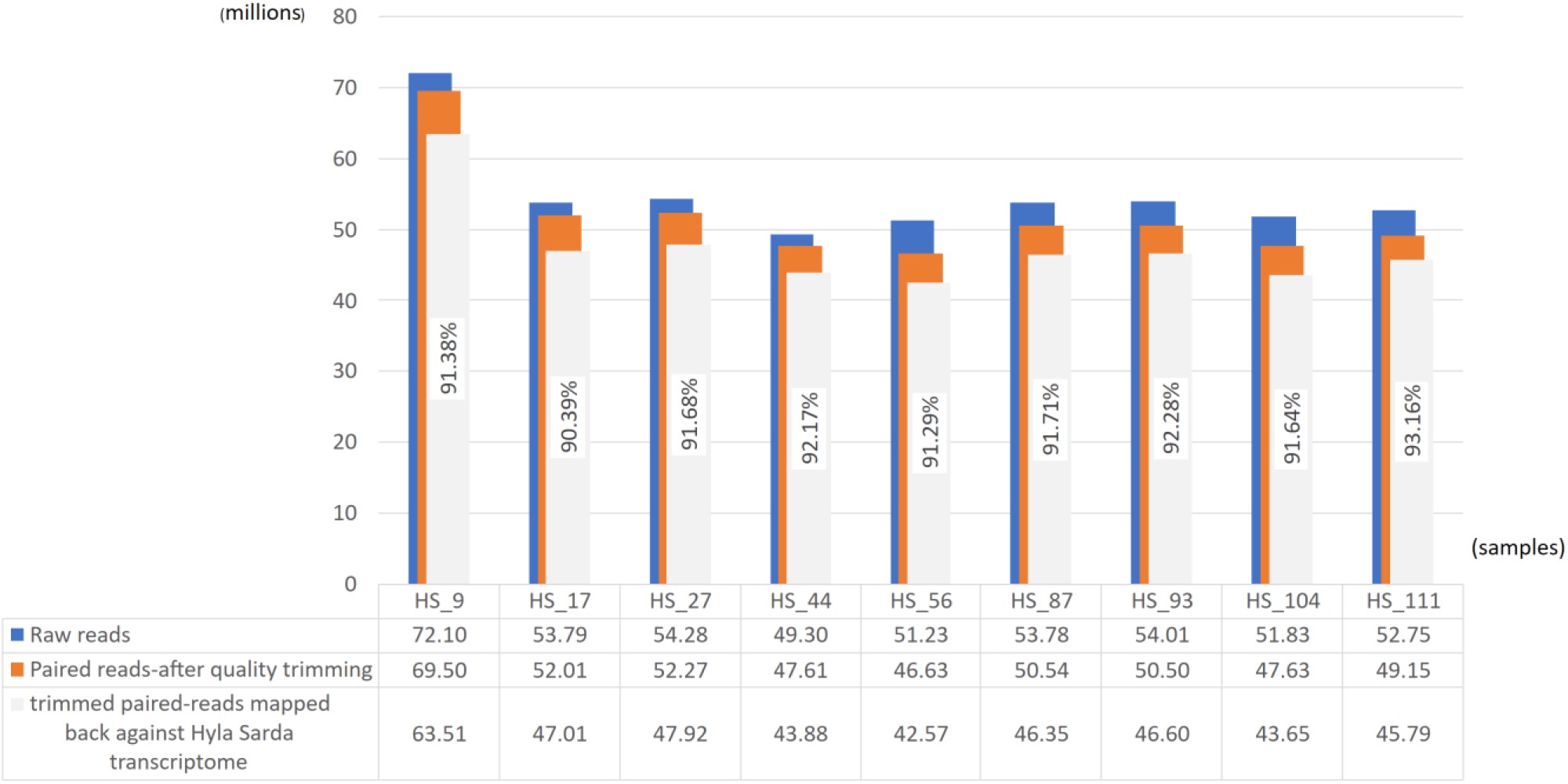
Paired-reads mapped back against *Hyla sarda* de-novo transcriptome. For each sample we have in blue the representation of total paired-reads, in orange the total paired-reads after the adapter removal and quality trimming and in grey we have the trimmed paired-reads mapped mapped-back against the *Hyla sarda* assembled de novo transcriptome.

TransRate assessment showed high values for the “TransRate Optimal Score” item, remaining around 0.09. The value of “good contigs” decreased after CD-HIT-est (due to redundancy removing), but with a value of 0.85 for the final assembly. TransRate also reported a value of GC around 44%. The final assembled transcriptome included a total of 1295741 transcripts with an N50 of 914 bp.

The BUSCO assessment also showed a high value for the “Complete and single-copy” equal to 124 (48.6 %) after the CD-HIT-est processing. As shown in Supplementary Table 3, CD-HIT-est improved the assembled transcriptome (“Complete and single-copy” higher than single rnaSPAdes and Trinity de-novo assemblies, Supplementary Table 3) removing the redundancy, reducing the number of transcripts generated by the 2 assemblers, improving the quality scores.

The transcriptome was functionally annotated by performing DIAMOND and InterProScan. By selecting the best hit for each database, the annotation matrix generated with Diamond has led to 88671 and 132635 contigs annotated in at least one database.

InterProScan is a tool that combines different protein signature recognition methods of the InterPro member databases into one resource. It provides as result the corresponding InterPro accession numbers and, among other accession IDs, the GO and KEGG annotation.

#### Code availability

All the software programs used in this article (de-novo transcriptome assembly, pre-and post-assembly steps and transcriptome annotation) are listed with the version in the Methods paragraph. In case of no details on parameters, the programs were used with the default settings.

## Data Records

All the datasets generated in this project were deposited in the European Nucleotide Archive (ENA) database under project identification number PRJEB51203 (ENA accessions are listed in table 1). Datasets containing the raw Trinity and rnaSPAdes transcriptome assemblies, unigenes, and functional annotation files were deposited on *figshare* archive (https://figshare.com/; links to pipeline outcomes are listed in Supplementary Table 4).

## Data Availability Statement

The datasets presented in this study can be found in online repositories. The names of the repository/repositories and accession number(s) can be found in the article/Supplementary Material.

## Author Contributions

DC conceived and financed the study; DC, AC and RB designed the experiment; AC, RB and GS performed sample collection and preparation; AC coordinated the RNA extraction and sequencing; TC designed and coordinated the bioinformatic analysis; PL and TC performed reads quality assessment, reads alignment on transcriptome, transcriptome annotation and validation; PL, TC, RB and AC, drafted the manuscript; All authors reviewed and approved the manuscript.

## Acknowledgments

We are grateful to Michela Paoletti for her support during the laboratory procedures, and to Jessica Di Martino for her support in bioinformatics analysis.

## Supplementary Material

The Supplementary Material can be found online at [to be populated after acceptance].

