## Supplementary material for "First brain de-novo transcriptome of Tyrrenhian tree frog, *Hyla sarda*, for the study of dispersal-related behavioral variation"

### Supplementary Figures and Tables

**Supplementary Figure 1.** The cleaned reads from all samples were assessed with FastQC and visualized with MultiQC. (a) Read count distribution for mean sequence quality. (b) Mean quality scores distribution. (c) Read length distribution. (d) Mean quality scores distribution.

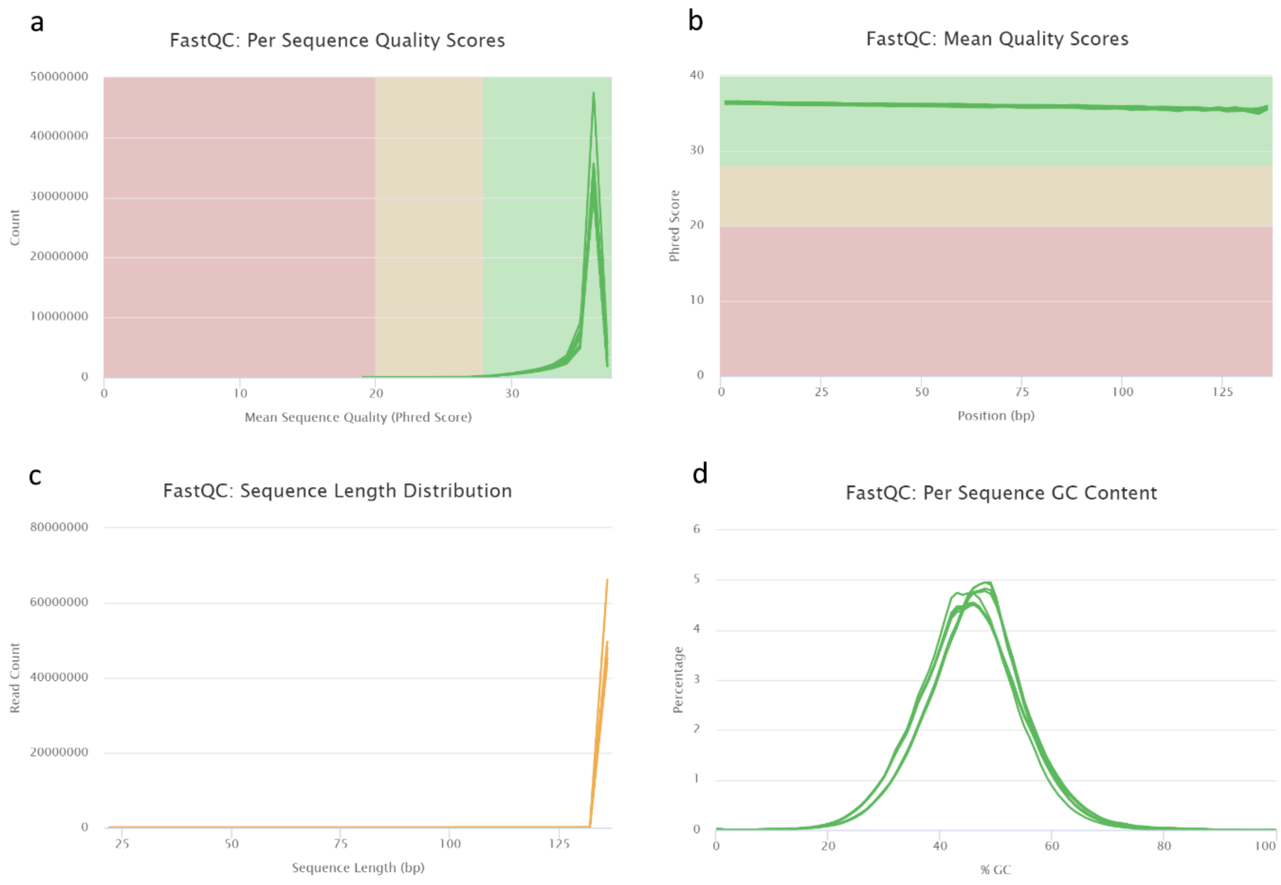

**Supplementary Figure 2.** Workflow of the bioinformatic pipeline, from raw data to annotated scripts, for the de novo transcriptome assembly of *Hyla sarda*.

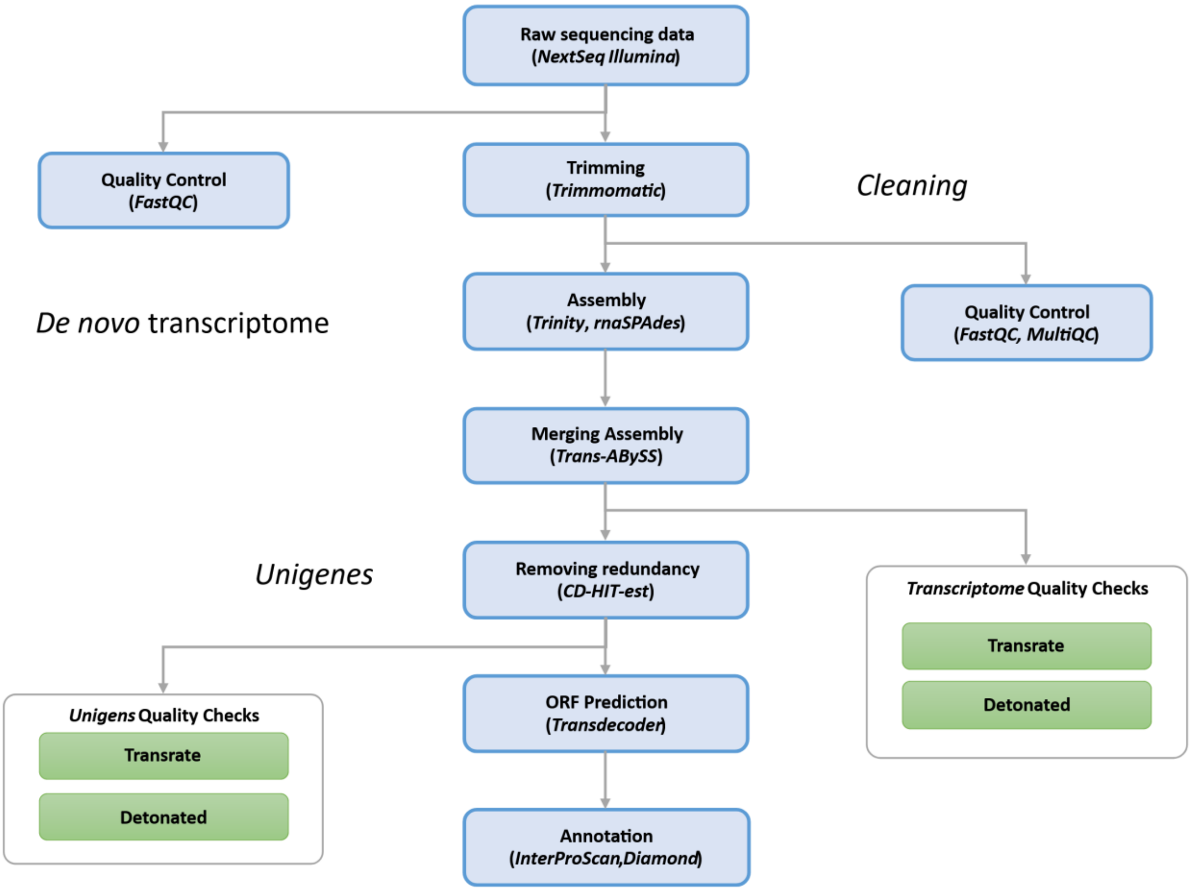

**Supplementary Figure 3.** Venn diagrams for the number of contigs annotated with Diamond (Blastx (a) and Blastp (b) functions) against the three databases:  NCBI nr, Swiss-Prot, TrEMBL.

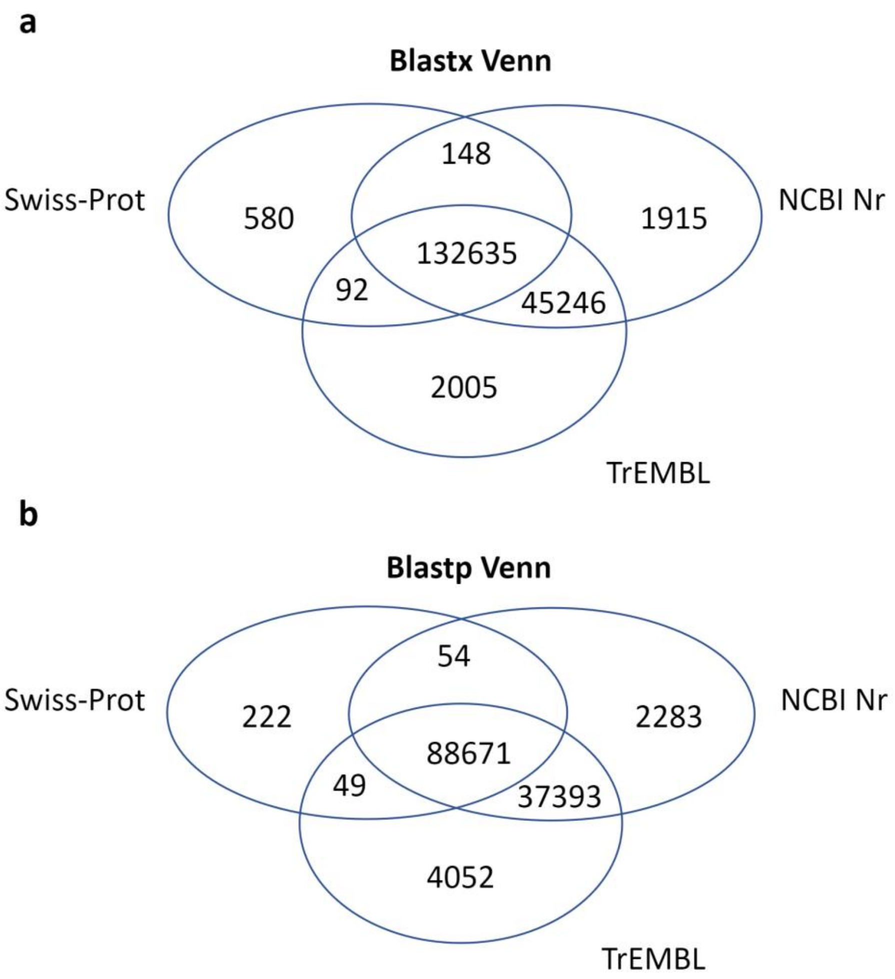

**Supplementary Table 1.** Statistics on rnaSPAdes and Trinity output and the result after CD-HIT-est (Unigenes).

| | Trinity | rnaSPAdes | CD-HIT-est (Unigenes) |
|---|---|---|---|
| **TRANSRATE v.1.0.3** | | | |
| Total transcripts | 1580186 | 1244846 | 1295741 |
| N50 | 684 | 1475 | 914 |
| GC content (%) | 44.00 | 44.00 | 44.00 |
| TransRate Assembly Score | 0.0277 | 0.0288 | 0.0519 |
| TransRate Optimal Score | 0.0936 | 0.0459 | 0.0933 |
| TransRate Optimal Cutoff | 0.3149 | 0.0155 | 0.0144 |
| p good contigs | 0.48 | 0.88 | 0.85 |
| **BUSCO v.3.0.2** | | | |
| Complete BUSCOs (C) | 253 (99.2%) | 247 (96.8 %) | 251 (98.4%) |
| Complete and single-copy BUSCOs (S) | 80 (31.4 %) | 97 (38.0 %) | 124 (48.6%) |
| Complete and duplicated BUSCOs (D) | 173 (67.8 %) | 150 (38.8 %) | 127 (49.8 %) |
| Fragmented BUSCOs (F) | 1 (0.4 %) | 5 (2.0 %) | 2 (0.8 %) |
| Missing BUSCOs (M) | 1 (0.4%) | 3 (1.2 %) | 2 (0.8 %) |
| Total BUSCO groups searched | 255 | 255 | 255 |

**Supplementary Table 2**. Summary of annotations on different databases.

| Annotation statistics | |
|---|---|
| **Number of blastx results** | |
| NCBI nr | 179944 (46.99 %) |
| Swiss-Prot | 133455 (34.85 %) |
| TrEMBL | 179978 (47.00 %) |
| **Number of blastp results** | |
| NCBI nr | 128401 (33.53 %) |
| Swiss-Prot | 88996 (23.24 %) |
| TrEMBL | 130165 (33.99 %) |

**Supplementary Table 4.** Overview of data files/data sets.

| Label | Name of data file/data set | File types (file extension) | Data repository and identifier (DOI or accession number) |
|---|---|---|---|
| Data file 1 | Trinity RNA-Seq de novo transcriptome assembly | Fasta file (.fa) | https://figshare.com/articles/online_resource/Assembly_HS_/17005156?file=31457608 |
| Data file 2 | rnaSPADES RNA-Seq de novo transcriptome assembly | Fasta file (.fa) | https://figshare.com/articles/online_resource/Assembly_HS_/17005156?file=31457605 |
| Data file 3 | Hyla Sarda RNA-Seq de novo transcriptome assembly (Unigenes) | Fasta file (.fa) | https://figshare.com/articles/online_resource/Assembly_HS_/17005156?file=31457614 |
| Data file 4 | Open reading frames (ORFs) prediction | Fasta file (.fa) | https://figshare.com/articles/online_resource/Assembly_HS_/17005156?file=31457617 |
| Data file 5 | Functional annotation from non-redundant (nr) NCBI | Text file (.txt) | https://figshare.com/articles/online_resource/Annotation_HS_/17005198?file=31457620 |
| Data file 6 | Functional annotation from Swiss-Prot | Text file (.txt) | https://figshare.com/articles/online_resource/Annotation_HS_/17005198?file=31457623 |
| Data file 7 | Functional annotation from TrEMBL UniProt | Text file (.txt) | https://figshare.com/articles/online_resource/Annotation_HS_/17005198?file=31457632 |
| Data file 8 | Functional annotation from non-redundant (nr) protein NCBI | Text file (.txt) | https://figshare.com/articles/online_resource/Annotation_HS_/17005198?file=31457635 |
| Data file 9 | Functional annotation from Swiss-Prot Protein | Text file (.txt) | https://figshare.com/articles/online_resource/Annotation_HS_/17005198?file=31457626 |
| Data file 10 | Functional annotation from TrEMBL UniProt Protein | Text file (.txt) | https://figshare.com/articles/online_resource/Annotation_HS_/17005198?file=31457629 |
| Data file 11 | InterProScan results | Text file (.txt) | ??? |
